# Prohaptoglobin Promotes Pancreatic Cancer Progression by sustaining YAP Activity

**DOI:** 10.64898/2026.07.26.740853

**Authors:** Jumpei Kondo, Honoka Nakayama, Ayusa Kuroda, Ayumu Hayashibara, Daisuke Sakon, Shinji Takamatsu, Hirofumi Akita, Hidetoshi Eguchi, Eiji Miyoshi

## Abstract

**Background & Aims:** Prohaptoglobin (proHp), a precursor of haptoglobin (Hp), has recently emerged as a cancer-associated biomarker, but its functional role in pancreatic cancer remains unclear. We investigated whether proHp promotes malignant phenotypes of pancreatic cancer and explored signaling pathways involved.

**Methods:** Serum proHp was examined in patients with pancreatic cancer and healthy controls. *HP* expression in pancreatic tumors and cell lines was analyzed using transcriptomic datasets. ProHp function was evaluated using PSN1 *HP* knockout (HPKO) cells, exogenous proHp supplementation, xenograft models, and RNA sequencing of PSN1 wild-type and HPKO cells. YAP activity was assessed by target gene expression, subcellular localization, YAP overexpression, and inhibition of YAP–TEAD interaction.

**Results:** Serum proHp was significantly elevated in patients with pancreatic cancer, and a subset of tumors and pancreatic cancer cell lines showed *HP* expression comparable to liver, indicating a tumor-derived source of proHp. In PSN1 cells, HPKO reduced motility, whereas exogenous proHp partially rescued this defect and enhanced motility in additional pancreatic cancer cell lines. Wild-type PSN1 cells continued to proliferate beyond confluence and formed rapidly growing xenograft tumors, which were abolished in HPKO cells. At high density, wild-type cells maintained YAP-related gene expression and nuclear YAP despite Hippo activation, whereas HPKO exhibited reduced nuclear YAP, indicating noncanonical YAP regulation by proHp. YAP restoration in HPKO cells rescued high-density proliferation and cell motility, while a YAP–TEAD inhibitor selectively reduced high-density proliferation of wild-type but not HPKO cells.

**Conclusion:** ProHp promotes pancreatic cancer progression in a context-dependent manner by sustaining YAP activity and enabling cells to partially overcome contact-dependent growth inhibition.

**Synopsis:** Prohaptoglobin, a precursor of haptoglobin, is elevated in pancreatic cancer and produced by tumor cells. It promotes cell motility, supports tumor growth under high-density conditions, and maintains YAP–dependent transcription that overrides contact-dependent growth inhibition.

**What You Need to Know:** 

**Background:** Prohaptoglobin, a precursor of haptoglobin, is elevated in pancreatic cancer, but its tumor-derived origin and functional role are unknown. We examined whether prohaptoglobin drives progression by sustaining YAP signaling.

**Impact:** We show that tumor-derived prohaptoglobin sustains nuclear YAP activity, allowing pancreatic cancer cells to bypass contact inhibition and proliferate, revealing prohaptoglobin as a context-dependent driver rather than a passive biomarker.

**Future Directions:** Defining how prohaptoglobin engages upstream Hippo–YAP regulators and whether prohaptoglobin–YAP signaling is targetable in vivo may uncover new biomarkers and therapeutic vulnerabilities for pancreatic cancer.

## Introduction

Haptoglobin (Hp) is a liver-derived acute-phase protein that exerts multiple functions in the context of inflammation and tissue injury. Among these, the best-characterized role is its function as a high-affinity scavenger of free hemoglobin (Hb) released during hemolysis, forming Hp–Hb complexes that are subsequently cleared via CD163-expressing macrophages and thereby limiting Hb/heme-induced oxidative tissue damage ^1^. Consistent with this protective scavenging function, Hp expression is markedly upregulated during systemic inflammation and tissue injury as part of the hepatic acute-phase response. Haptoglobin also exerts immunomodulatory functions, largely through Hp–Hb uptake by CD163-expressing macrophages, which alters macrophage activation states and cytokine profiles ^2^. For example, this process can influence the balance between Th1- and Th2-type adaptive immune responses ^3^. In healthy individuals, serum levels of fucosylated Hp (Fuc-Hp) are low, whereas various cancers and liver diseases are associated with increased core fucosylation ^4^ and aberrant release of Fuc-Hp into the circulation ^5^. In patients with cancer, elevated serum Fuc-Hp levels have been reported in several tumor types, including pancreatic ^6, 7^, colorectal ^8^, prostate ^9^, ovarian ^10^, and lung cancers ^11^, as well as hepatocellular carcinoma ^12^. We also previously reported that Fuc-Hp levels increase in parallel with clinical stage progression in pancreatic cancer ^13^.

Traditionally, Fuc-Hp has been detected indirectly using sandwich ELISA with anti-haptoglobin antibody and fucose-binding lectin such as AAL and PhoSL. We previously established and reported a monoclonal antibody, 10-7G, which directly recognizes Fuc-Hp ^13^. Notably, this 10-7G antibody also recognizes prohaptoglobin (proHp), the precursor of mature Hp ^14^. ProHp is cleaved by the liver-enriched protease C1RL, yielding the α- and β-chains that form mature Hp linked by disulfide bonds ^15^. Although proHp is a transient precursor molecule under physiological conditions, several studies have shown that it can be aberrantly secreted in pathological states, including ovarian, prostate, and colorectal cancers ^14, 16, 17^. In addition, proHp possesses distinct biological functions not observed in mature Hp, and has been reported to be involved in inflammation and angiogenesis ^18, 19^. We reported that in colorectal cancer, proHp is elevated in association with disease stage and functions as a biomarker of progression, while promoting cell motility through epithelial–mesenchymal transition (EMT)-like changes ^14^. These observations suggest that proHp is not merely a passive precursor of Hp, but an active modulator of the tumor microenvironment. However, the mechanisms underlying the cancer-promoting functions of proHp remain poorly understood, and the role of proHp in pancreatic cancer has not yet been clarified.

In this study, we investigated the clinical significance and biological functions of proHp in pancreatic cancer, and sought to identify downstream signaling pathways that mediate its pro-tumorigenic effects.

## Materials and Methods

### Human samples and Animal studies

Serum samples were obtained from 64 patients with pancreatic cancer who underwent resection surgery at The University of Osaka Hospital and 64 healthy individuals who underwent medical checkup at aMs New Otani Clinic. This study was approved by The University of Osaka Hospital Ethical Review Board (approval number: 13563) and conducted in accordance with the Declaration of Helsinki. Animal studies were approved by the Institutional Animal Care and Use Committee of The University of Osaka Graduate School of Medicine, Division of Health Sciences (approval number: R5-03-01) and were performed in accordance with institutional and national guidelines for the care and use of laboratory animals.

### Cell Culture

PSN1, BxPC-3, PCI6 and HEK293T cells were obtained from American Type Culture Collection (ATCC, Virginia, USA) and PANC-1, PK-8, PK-45 and MIAPaCa-2 cells were purchased from RIKEN BioResource Research Center (Saitama, Japan). PSN1, PANC-1, BxPC-3, PCI6, and PK-8 cells were cultured in RPMI1640 (Nacalai Tesque, Kyoto, Japan). MIAPaCa-2 cells were cultured in Dulbecco’s modified Eagle’s medium (DMEM) low-glucose (Nacalai Tesque). HEK293T cells were cultured in Dulbecco’s modified Eagle’s medium (DMEM) high-glucose (Nacalai Tesque). The culture media for all cell lines were supplemented with 10% fetal bovine serum (FBS), 100 U/mL penicillin, and 100 µg/mL streptomycin and cultured at 37°C in a humidified atmosphere containing 5% CO2. Cell proliferation was evaluated either by counting cell number or by ATP assay using CellTiter-Glo (Promega, Madison, WI) according to the manufacturer’s instructions, in which luminescence was measured with an INFINITE M PLEX plate reader (Tecan, Männedorf, Switzerland).

### Establishment of genetically modified cell lines

PSN1 cells were genetically modified using a lentiviral vector system to generate *HP* knockout and YAP-overexpressing cell lines. The pLentiCRISPR v2 vector (Addgene #52961) was used for *HP* knockout, while pLX304-YAP (Addgene #42555) and pLX304-YAP(S6A) (Addgene #42562) were used for YAP overexpression. All vectors, except for the *HP*-targeting construct, were obtained from Addgene (Watertown, MA, USA). For generation of the *HP* knockout construct, the pLentiCRISPR v2 vector was digested with BsmBI and ligated with annealed guide RNA oligonucleotides targeting exon 2 of *HP* (5′-GAGGGCAATGACAGCTCCCA-3′). Lentiviral particles were produced in HEK293T cells by co-transfection of each expression vector with pMD2.G (Addgene #12259) and psPAX2 (Addgene #12260). Viral supernatants were collected and used to transduce PSN1 cells in the presence of polybrene (4 µg/mL), followed by spin infection (2000 rpm, room temperature, 60 min). Stable cell lines were selected with puromycin (2 µg/mL) for *HP* knockout or blasticidin (2 µg/mL) for YAP overexpression. For establishment of proHp-overexpressing PANC1 cells, cells were transfected with a proHp expression vector in pcDNA3.1 backbone^14^ using polyethyleneimine (PEI), followed by selection with hygromycin B (200 µg/mL). Knockout and overexpression were confirmed by western blotting.

### Purification of proHp

Purified proHp was prepared as previously described ^14^. Briefly, the supernatant from proHp-FLAG–overexpressing HEK293T cells was collected, and proHp-FLAG was purified by affinity chromatography using ANTI-FLAG M2 Affinity Gel (Sigma-Aldrich, St. Louis, MO, USA) according to the manufacturer’s instructions.

### Wound Repair Assay

The protocol used for the wound repair assay was described previously ^14^. Briefly, cells were seeded in 6-well plates and cultured to confluence. A P1000 pipette tip was used to make a linear scratch in the cell monolayer. The culture medium was then removed and replaced with fresh RPMI 1640 medium containing 2% FBS and 5 µg/mL mitomycin C (Nacalai Tesque), and the plates were incubated at 37°C in a humidified atmosphere of 5% CO₂. For the proHp rescue conditions, 20µg/mL of proHp was added to the culture medium. Phase-contrast images were acquired using a BZ-9000 fluorescence microscope (Keyence, Osaka, Japan) immediately after scratching and 24 h later. The wound area was quantified using ImageJ software (NIH, Bethesda, MD).

### Xenograft tumor formation assay

Six-week-old BALB/c-nu and NOD/SCID mice (Jackson Laboratory Japan, Kanagawa, Japan) were inoculated subcutaneously with 4.0 x 10^5^ PSN1 WT or HPKO cells suspended in 100 µL of phosphate-buffered saline (PBS), mixed with an equal volume of Matrigel (Corning, Corning, NY), using a 27-gauge needle. Tumor volume and body weight were measured at regular intervals. Tumor volume was calculated using the formula: volume = (width)^2^ × length / 2. Mice were euthanized for ethical reasons when tumor volume exceeded 2,000 mm³, in accordance with institutional animal care guidelines. Xenograft tumors were either processed as formalin-fixed, paraffin-embedded (FFPE) specimens for hematoxylin and eosin (H&E) staining or embedded directly in OCT compound (Sakura Finetek, Tokyo, Japan) for preparation of frozen tissue sections.

### Western blot

Cells were lysed in RIPA buffer (50 mM Tris-HCl, pH 7.6, 150 mM NaCl, 1% Nonidet P-40, 0.5% sodium deoxycholate, 0.1% SDS) supplemented with protease inhibitor cocktail (Nacalai Tesque) and phosphatase inhibitor cocktails I and II (Sigma-Aldrich). Proteins were electrophoresed on polyacrylamide gels and transferred onto PVDF membranes (Merck Millipore). After blocking with Tris-buffered saline containing 0.1% Tween 20 (TBST) and 5% skim milk, proteins were probed with the following primary and secondary antibodies listed in the Supplementary Table S1. Chemiluminescent signals were developed using Chemi-Lumi One Super (Nacalai Tesque) and detected with a FUSION chemiluminescence imaging system (Vilber Lourmat, Collégien, France).

### Semi-quantitative real time RT-PCR

Total RNA was extracted using RNeasy Mini Kit (QIAGEN, Hilden, Germany) according to the manufacturer’s instructions. cDNA was synthesized using GoScript Reverse Transcription System (Promega) in a T100 Thermal Cycler (Bio-Rad, Hercules, CA). A 10-fold dilution of the synthesized cDNA was used for semi-quantitative real-time reverse transcription PCR (RT-PCR) with THUNDERBIRD SYBR qPCR Mix containing ROX reference dye (Toyobo, Osaka, Japan) on a StepOnePlus Real-Time PCR System (Thermo Fisher Scientific, Waltham, MA). Relative mRNA expression levels were calculated by the ΔΔCt method using GAPDH as the reference gene. The primer pairs used are listed in Supplementary Table S2.

### Immunofluorescence imaging

For immunocytochemistry, cells were fixed with 4% paraformaldehyde (PFA) on ice for 15 min. For immunofluorescence imaging of tissue sections, samples were fixed with 10% neutral buffered formalin (NBF) for 10 min at room temperature. The samples were then washed, and blocked with 2.5% goat serum in PBS, and incubated with rabbit anti-YAP antibody (Cell Signaling Technology) followed by Alexa Fluor 488–conjugated anti-rabbit IgG secondary antibody (Thermo Fisher Scientific) and counterstained with DAPI (Nacalai Tesque). Fluorescence images were acquired using a BZ-9000 fluorescence microscope (Keyence, Osaka, Japan).

### RNA sequencing (RNA-seq) analysis

Total RNA was isolated from PSN1 WT and HPKO cells, as described in the section “Semi-quantitative real time RT-PCR”. The library was prepared using the TruSeq stranded mRNA sample prep kit (Illumina, San Diego, CA, USA) according to the manufacturer’s instructions and sequenced on an Illumina NovaSeq 6000 platform in the 101 bp single-end mode. Sequenced reads were QCed with trimmomatic v0.38, mapped to the human reference genome sequence using TopHat v2.1.1 in conjunction with Bowtie2 ver. 2.3.5.1 and SAMtools ver. 1.2. Fragment counts per kilobase per exon (FPKMs) were calculated using Cufflinks ver. 2.2.1.

The following RNA-seq data analysis was performed using R (versions 4.3.2–4.5.2). Differential gene expression (DGE) analysis was conducted using the edgeR ^20^ and DESeq2 ^21^ packages. Volcano plots were generated using the EnhancedVolcano package (https://github.com/kevinblighe/EnhancedVolcano). Clustering analysis and heatmap construction were performed using the heatmap3 package ^22^. Additional data visualization and graphical representations were created using ggplot2 ^23^.

Gene Ontology (GO) analysis was performed on the extracted gene lists using the ToppFun function in the ToppGene Suite ^24^. Gene set enrichment analysis (GSEA) was performed using GSEA v4.4.0 ^25^. FDR q value 0.1 was used as cutoff. The gene sets used for each enrichment analysis are specified in the Supplementary Table S3 (ref) ^26–28^. The data have been deposited in the DNA Data Bank of Japan (DDBJ) Sequence Read Archive (DRA) under accession number DRA028777.

### Database Analysis

Gene expression analysis of HP in normal pancreatic tissue, pancreatic cancer tissue, and normal liver tissue was performed using GENT2 ^29^ with data from the GPL570 platform. Gene expression analysis of HP in non-cancerous cell lines, pancreatic cancer cell lines, and hepatocellular carcinoma cell lines was conducted using the DepMap Portal ^30^ with dataset version 24Q2.

### Statistical Analysis

Significance was tested using an unpaired Student’s t-test or Wilcoxon rank-sum test for single comparisons. Repeated-measures ANOVA was employed to compare tumor growth and cell proliferation trajectories between WT and KO groups, with all individual measurements analyzed across the study period. Multiple testing corrections were applied using the Bonferroni method for both tests. A P-value of < 0.05 was considered statistically significant.

## Results

### Prohaptoglobin is upregulated in pancreatic cancer

We have previously established the 10-7G monoclonal antibody 10-7G, which directly recognizes fucosylated haptoglobin (Fuc-Hp) as well as prohaptoglobin (proHp), the precursor of mature Hp. Using a 10-7G-based ELISA based, we reported that elevated serum 10-7G-reactive antigens are elevated in patients with pancreatic cancer ^13^. However, this assay could not distinguish proHp from mature Fuc-Hp. Because proHp is cleaved by the liver-enriched serine protease C1RL into the α- and β-chains that form mature Hp linked by disulfide bonds^15^, proHp and mature Fuc-Hp can be separated by molecular size under reducing conditions. We therefore performed semi-quantitative western blotting (WB) of serum samples from patients with pancreatic cancer and healthy individuals, and found a significant increase in proHp in sera of patients with pancreatic cancer (Fig. 1A, B).

**Figure. 1.**
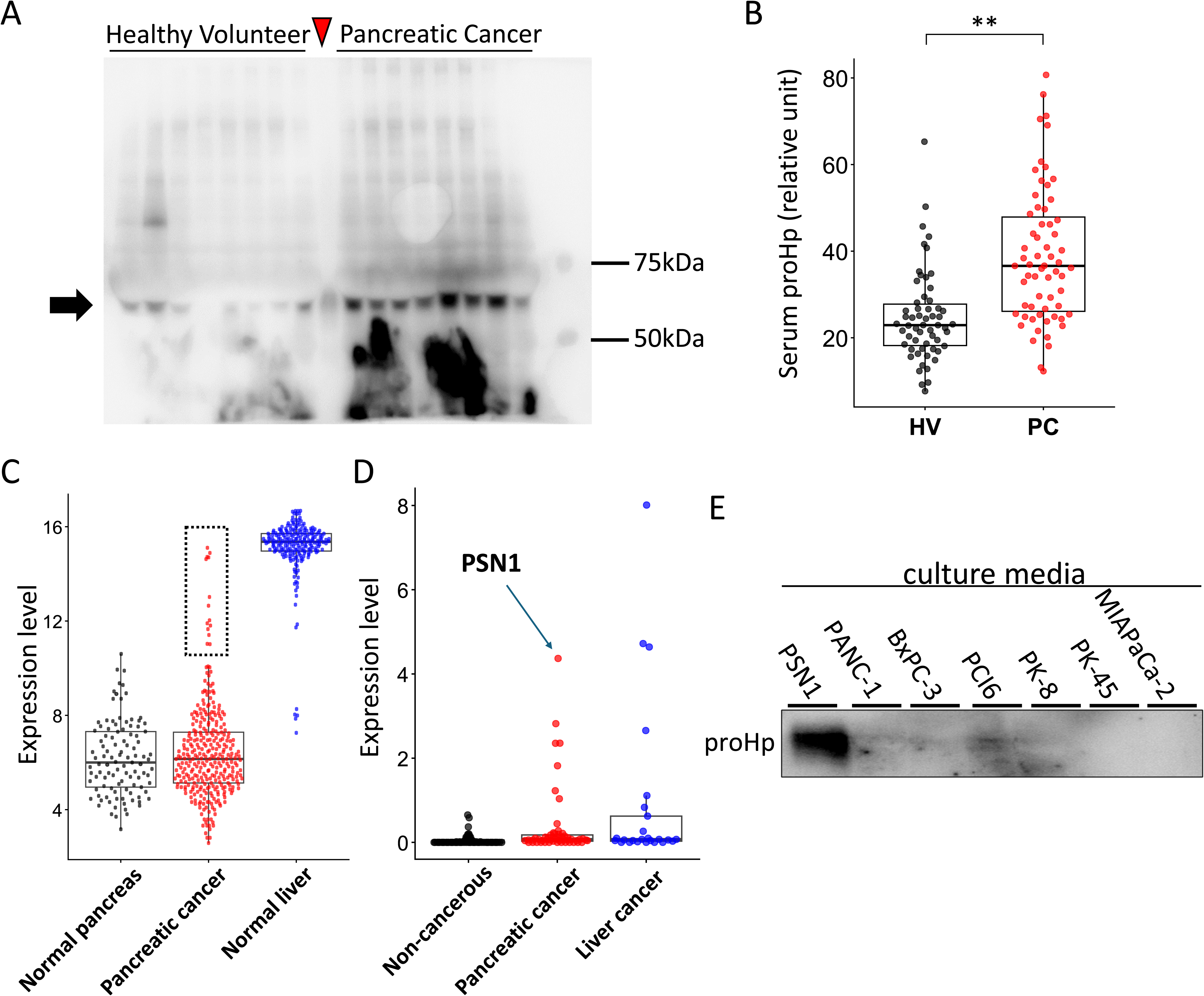
Prohaptoglobin is upregulated in pancreatic cancer. A. Representative immunoblot of sera from patients with pancreatic cancer and healthy volunteers labeled by the 10-7G antibody. Black arrow indicates the band size corresponding to proHp. Red arrowhead indicates the lane loaded 5µg of purified proHp. B. Box-and-Whisker plot of semi-quantified serum proHp levels in patients with pancreatic cancer (PC, N = 58) and healthy volunteers (HV, N = 67). **; P<0.01, Wilcoxon rank sum test. C. Box-and-Whisker plots of *HP* gene expression levels (log2 fold change) in normal pancreas (N = 105), pancreatic cancer (N = 324), and normal liver tissues (N = 215) from the GENT2 database. The dotted box indicates upper outliers in the pancreatic cancer group (21 tissues, 6.5%). D. Box-and-Whisker plots of *HP* gene expression levels (log2(TPM+1)) in cell lines of non-cancerous (N = 80), pancreatic cancer (N = 52), and liver cancer (N = 25) lineages from the DepMap database. E. Immunoblot of culture supernatants from pancreatic cancer cell lines labeled by the 10-7G antibody. The band size corresponding to proHp was presented.

Serum proHp can originate from multiple sources. We previously showed that immune cells can produce proHp ^31^, whereas colorectal cancer cells can serve as an autocrine source of proHp ^14^. To assess whether pancreatic cancer tissue also contribute, we analyzed *HP* gene expression in publicly available datasets. Among 324 pancreatic cancer cases, 21 cases (6.5%) exhibited *HP* gene expression levels comparable to those of liver, the primary Hp-producing organ (Fig. 1C). Similarly, analysis of pancreatic cancer cell lines revealed high *HP* expression, defined as greater than the median expression in liver-derived cell lines, in 7 of 52 cell lines (13.5%) (Fig. 1D). This is consistent with our previous pathological observation of a pancreatic cancer case with strong HP immunostaining in tumor cells ^31^. Taken together, these findings suggest that, at least in a subset of cases, pancreatic cancer cells are exposed to elevated levels of proHp through autocrine, paracrine, or hematogenous delivery.

### Prohaptoglobin stimulates the migratory capacity of pancreatic cancer cell lines

Recent studies have suggested that proHp is not merely an immature precursor that leaks into the circulation, but instead exerts biological activities distinct from those of mature Hp ^14, 18, 19^. Thus, elevated proHp in pancreatic cancer may not only serve as a biomarker, but could also actively contribute to tumor pathogenesis. To investigate this possibility, we examined the effects of proHp on malignant phenotypes using pancreatic cancer cell lines. Among pancreatic cancer cell lines in the public dataset, PSN1 showed the highest *HP* expression, and secreted detectable readily detectable levels of proHp (Fig. 1D, E). We generated *HP*-knockout PSN1 cells (HPKO) using CRISPR/Cas9 (Fig. 2A) and first evaluated their migratory capacity. In wound-repair assays performed in the presence of mitomycin to suppress proliferation, HPKO cells exhibited significantly reduced motility compared with wild-type (WT) cells (Fig. 2B, C). Notably, exogenous addition of purified proHp partially rescued the migratory defect of HPKO cells in a statistically significant manner (Fig. 2B, C).

**Figure. 2.**
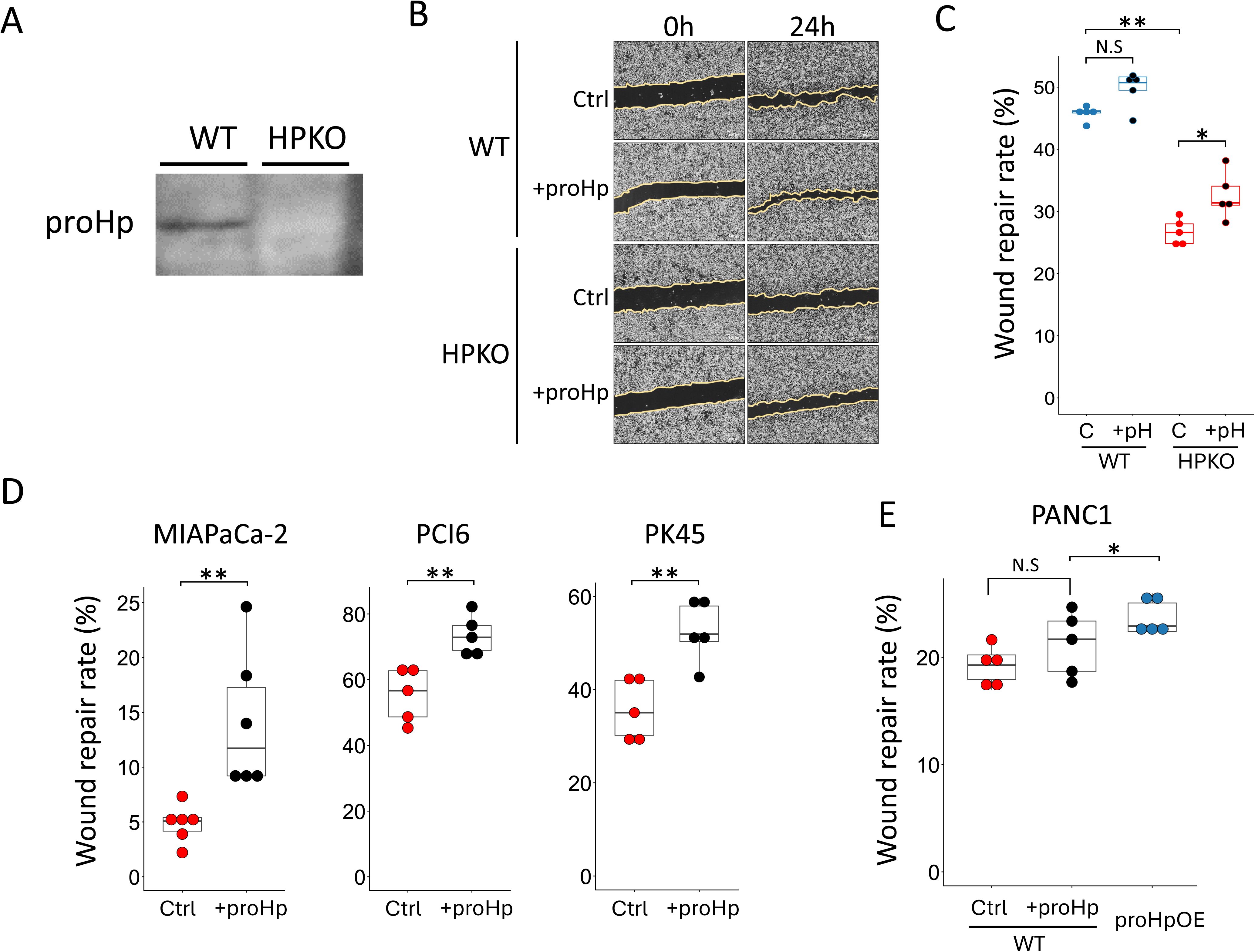
Prohaptoglobin stimulates the migratory capacity of pancreatic cancer cell lines. A. Immunoblot of culture supernatants from PSN1 WT and HPKO cell lines labeled by the 10-7G antibody. B. Representative phase-contrast microscopic images of wound repair assays for PSN1 WT and HPKO cells at 0 and 24 hours post-scratching, cultured without (Ctrl) or with supplementation of purified proHp (+proHp). C-E. Box-and-Whisker plots quantifying wound repair rates in PSN1 WT and HPKO (C), MIAPaCa-2, PCI6, and PK45 cells (D) and PANC1 WT and proHp-overexpressing (proHpOE) cells (E). Cells were cultured without (C/Ctrl) or with supplementation of purified proHp (+pH/+proHp). *; P<0.05, **; P<0.01, N.S; not significant, Wilcoxon rank sum test.

We next examined whether extracellular proHp can enhance motility in pancreatic cancer cell lines. Addition of purified proHp to the culture medium significantly accelerated wound closure in MIAPaCa-2, PCI6, and PK45 cells, all of which showed undetectable proHp production (Fig. 1E, 2D). In PANC1 cells, exogenous proHp tended to increase motility without reaching statistical significance, whereas forced HP overexpression significantly enhanced wound closure (Fig. 2E). These results indicate that proHp promotes the migratory capacity of pancreatic cancer cells as a secreted protein, acting either in an autocrine or a paracrine manner.

### Prohaptoglobin promotes pancreatic cancer cell growth in a context-dependent manner

We next investigated the impact of proHp on cell proliferation. In contrast to its clear effect on motility, *HP* knockout did not affect in vitro proliferation of PSN1 during the exponential growth phase up to confluence (Fig. 3A). Given this dissociation between enhanced motility and unchanged proliferation *in vitro*, we examined whether proHp contributes to pancreatic cancer progression *in vivo*. When PSN1 WT and HPKO cells were subcutaneously transplanted into nude mice, HPKO tumors exhibited significantly reduced growth compared with WT tumors (Fig. 3B, C). Similar growth suppression of HPKO tumors was observed in NOD/SCID mice, which lack both T and B cells (Fig. 3D), indicating that the growth disadvantage of HPKO cells is primarily attributable to tumor cell-intrinsic mechanisms rather than host adaptive immune responses.

**Figure. 3.**
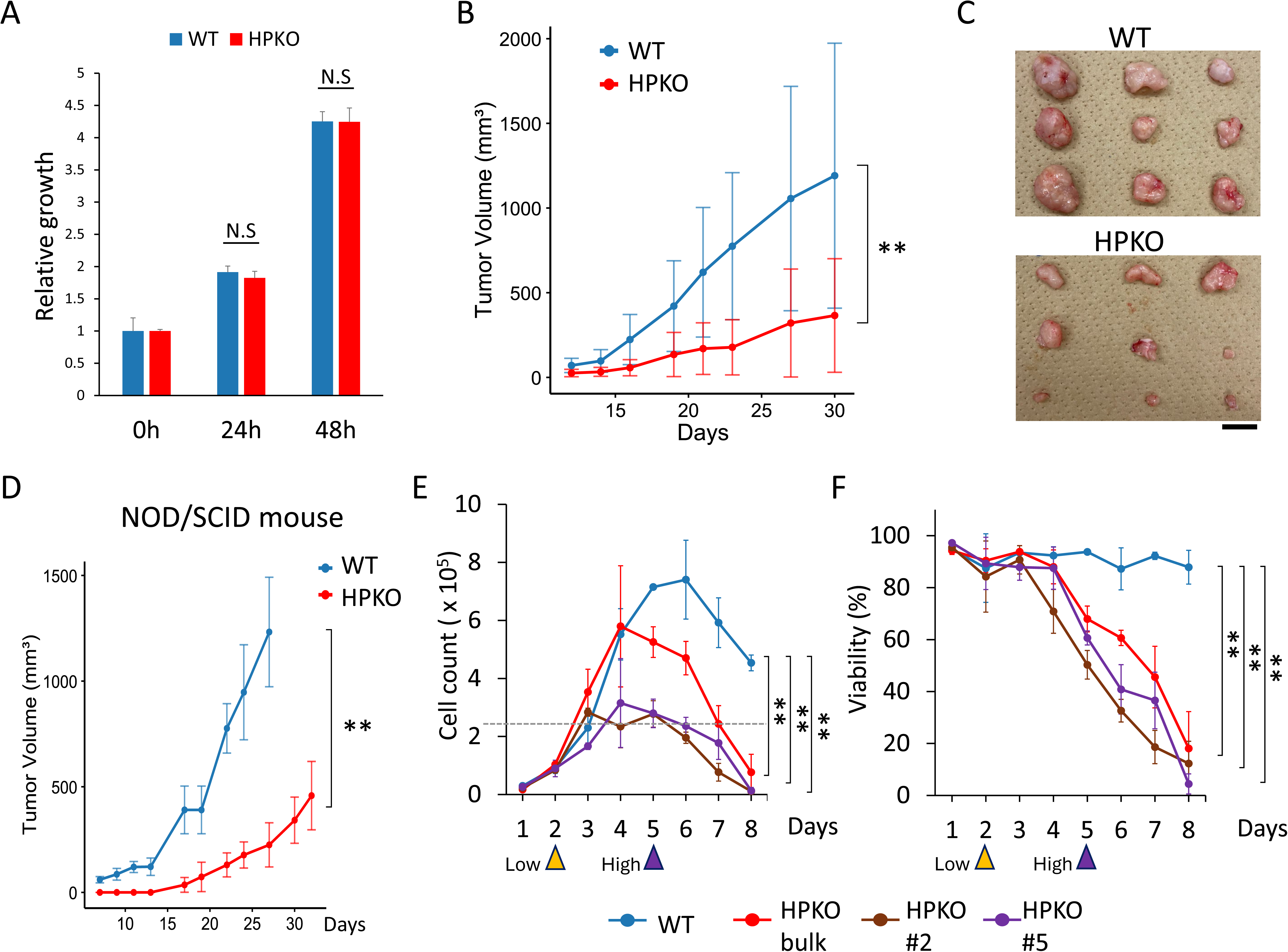
Prohaptoglobin promotes pancreatic cancer cell growth in a context-dependent manner. A. Cell growth assay of PSN1 WT and HPKO cells cultured for 24, and 48 hours. Viability was normalized to that at 0 hour. Data is shown as mean□+□SD. N.S; not significant, Student’s t-test with Bonferroni correction. B. Xenograft tumor growth curves of PSN1 WT (blue) and HPKO (red) cells subcutaneously transplanted into nude mice. N = 9 for each group, Data is shown as mean□±□SD. **: P<0.01, repeated-measures ANOVA. C. Macroscopic images of xenograft tumors derived from PSN1 WT (upper) and HPKO (lower) cells harvested at day 30 after transplantation. Scale bar, 1 cm. D. Xenograft tumor growth curves of PSN1 WT (blue) and HPKO (red) cells subcutaneously transplanted into NOD/SCID mice. N = 6 for each group, Data is shown as mean□±□SD. **: P<0.01, repeated-measures ANOVA. E-F. Growth curve (E) and cell viability (F) of PSN1 WT (blue), bulk HPKO (red), HPKO clone #2 (brown), and HPKO clone #5 (purple) for extended period. Horizontal dotted line in E denotes the cell density in which cells are visually confluent. Yellow and purple arrowheads in (E, F) indicate the time points at which cells were harvested under low-and high-density conditions, respectively, for subsequent RNA-seq and qPCR analyses shown in Figure 4. N = 3 for each condition, Data is shown as mean□±□SD. **: P<0.01, repeated-measures ANOVA.

To more comprehensively evaluate the proliferative potential of PSN1 cells, we next extended the *in vitro* culture period beyond visual confluence. WT cells continued to proliferate after reaching confluence, whereas HPKO cells ceased proliferation upon confluence and rapid lost viability thereafter (Fig. 3E, F). This phenotype was consistently observed not only in bulk HPKO cells selected by drug resistance but also in clonally derived HPKO#2 and HPKO#5 clones (Fig. 3E, F). This phenotype was also observed in PANC-1 cells, which produce low levels of proHp (Fig. 1E). Overexpression of proHp in PANC-1 cells promoted proliferation beyond confluence (Fig. S1). Collectively, these results demonstrate that proHp supports pancreatic cancer progression through tumor cell-intrinsic mechanisms. Importantly, the growth-suppressive phenotype of HPKO cells emerges only under high-density conditions, suggesting that proHp functions to override contact-dependent growth inhibition.

### proHp induces YAP-regulated gene expression

We next sought to elucidate the transcriptional programs regulated by proHp using RNA sequencing of PSN1 WT and HPKO cells cultured under low- or high-confluency conditions (Fig. S2A). Clustering analysis revealed that transcriptomic differences between WT and HPKO cells were more pronounced at high confluency than at low confluency (Fig. S2B). In line with the observation that WT and HPKO cells proliferated similarly at low density (Fig. 3A, E), we therefore focused subsequent analyses on differentially expressed genes (DEGs) under high-confluency conditions (Fig. 4A, B). Gene ontology analysis of genes downregulated in HPKO cells (i.e., proHp-induced genes) showed significant enrichment for categories related to cell signaling pathways (Fig. 4C, left). In contrast, genes upregulated in HPKO cells (i.e., proHp-repressed genes) were enriched for terms such as “inflammatory response” and “regulation of cell adhesion,” suggesting that proHp suppresses these programs under high-confluency conditions (Fig. 4C, right).

**Figure. 4.**
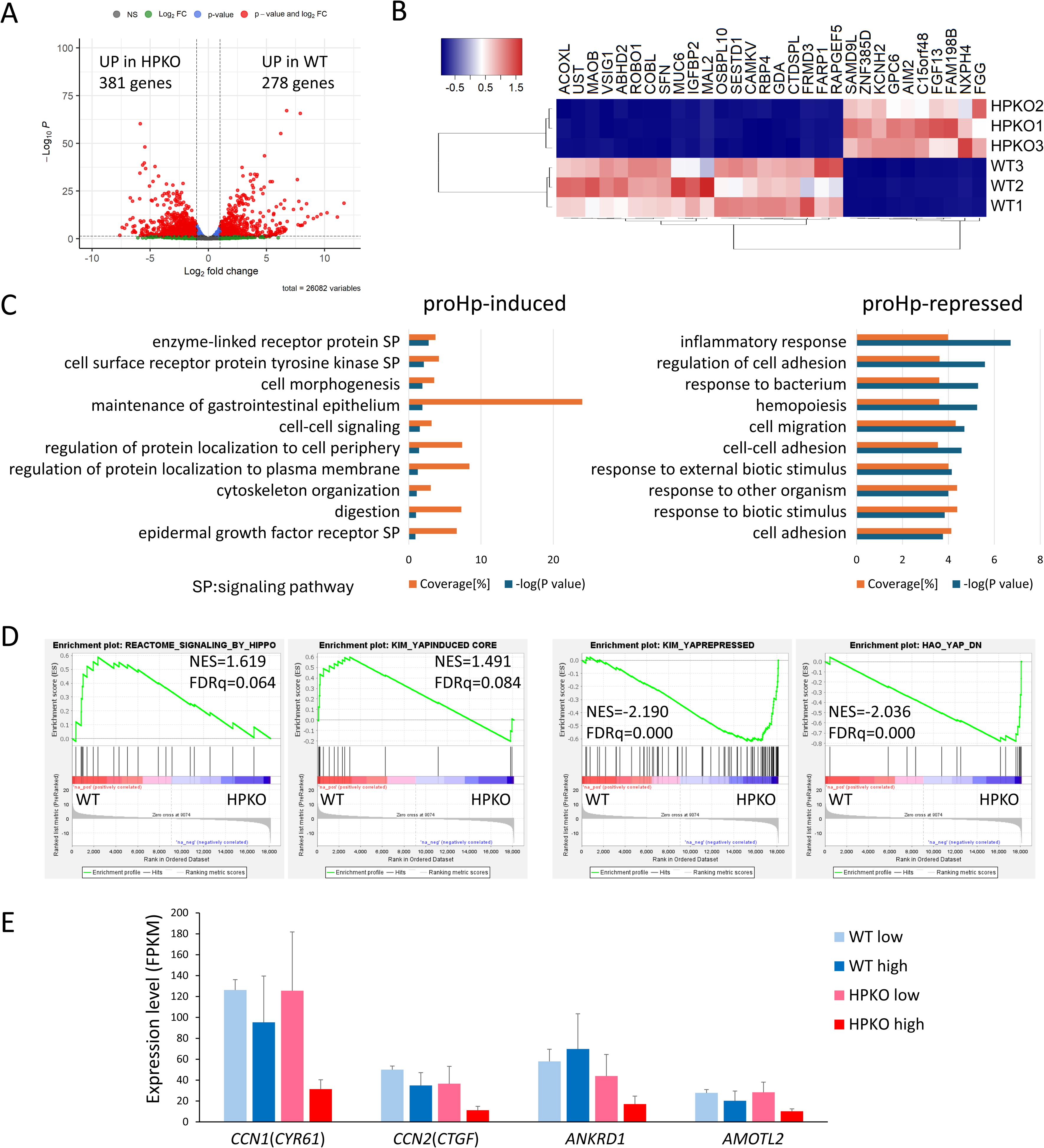
proHp induces YAP regulated gene expression in PSN1 cells. A. Volcano plot showing the differently expressed genes (DEGs) of PSN1 between WT and HPKO cells at high confluency. Red dots represent the genes with significantly altered expresseion level (>1.5-fold). B. Heatmap showing the top 30 DEGs ranked by the lowest P-values, comparing PSN1 WT and HPKO cells at high confluency. C. Gene ontology (GO) analysis of DEGs. 353 genes upregulated (>1.5-fold) in WT are defined as proHp-induced genes, and 374 genes upregulated (>1.5-fold) in HPKO are defined as proHp-repressed genes. Orange bars; coverage (percentage of input genes within each annotation), Blue bars: the -log10(1/P-value) for each GO term. D. Gene set enrichment analysis (GSEA) results comparing PSN1 WT and HPKO. Gene-set names are presented on top of each panel. Normalized Enrichment Score (NES) and False Discovery Rate q-value (FDRq) are indicated within each graph. E. Expression levels of selected YAP-target genes extracted from RNA-seq for PSN1 WT and HPKO cells cultured at low and high confluency. Data is shown as mean□+□SD.

Because Hippo signaling is a canonical pathway governing contact inhibition of proliferation ^32, 33^, we next examined the relationship between proHp status and Hippo pathway activity by Gene Set Enrichment Analysis (GSEA), focusing on Hippo signaling and its downstream co-activator YAP. In high-confluency cultures, WT cells showed enrichment of Hippo pathway and YAP target gene signatures, whereas HPKO cells showed enrichment of YAP-repressed gene sets (Fig. 4D). Consistently, the induction of representative YAP target genes, including *CCN1*(*CYR61*), *CCN2*(*CTGF*), *ANKRD1*, and *AMOTL2*, was attenuated in HPKO cells at high confluency, whereas WT cells maintained their expression (Fig. 4E). Together with the functional data, these findings support a model in which proHp prevents full activation of Hippo signaling at high cell density, thereby allowing YAP-dependent transcription and sustained proliferation despite increased cell–cell contact.

### proHp maintains YAP nuclear localization by limiting Hippo pathway activation

We next evaluated how proHp status affects YAP activation at high cell density in PSN1 cells. Consistent with the RNA-seq results, qPCR confirmed that expression of the YAP target genes *CCN1* and *CCN2* remained high in WT cells under high-confluency conditions in WT cells, whereas their expression was significantly reduced in HPKO cells (Fig. 5A). YAP functions as a transcriptional co-activator whose activity is negatively regulated by Hippo pathway-dependent phosphorylation at S127, which promotes its cytoplasmic retention and subsequent proteasomal degradation ^32, 33^. Interestingly, S127 phosphorylation of YAP was not increased in WT cells at high confluency (Fig. 5B, C). Independent of canonical Hippo pathway-dependent YAP regulation, noncanonical (i.e., Hippo-independent) mechanisms have been reported in which SRC family kinases (SFKs) promote YAP nuclear localization through tyrosine phosphorylation at Y357, regardless of S127 status ^34^. In PSN1 WT cells, but not in HPKO cells, high-confluency culture was associated with SFK activation and enhanced Y357 phosphorylation of YAP (Fig. 5B, C). Treatment of WT cells at high density with the SFK inhibitor dasatinib reduced Y357 phosphorylation while increasing S127 phosphorylation (Fig. 5D, E), indicating that Y357 phosphorylation under these conditions is SFK-dependent. Consistently, immunofluorescence analysis demonstrated prominent nuclear localization of YAP in WT cells under high-density conditions, whereas HPKO cells displayed a marked reduction in nuclear YAP (Fig. 5F, G, and S3). Of note, HPKO cells tended to occupy a larger projected cell area per cell under high confluency.

**Figure. 5.**
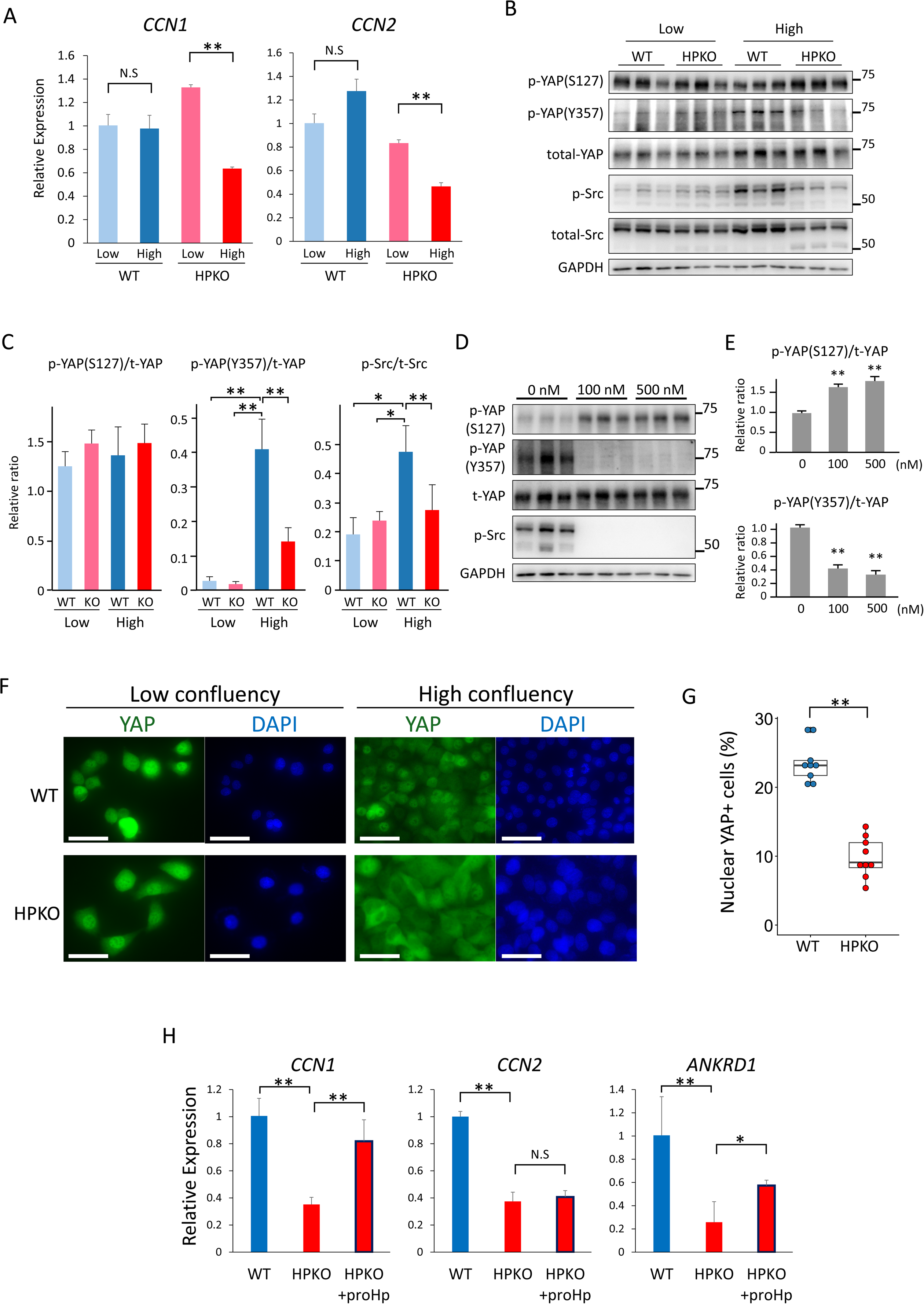
proHp maintains YAP nuclear localization by limiting Hippo pathway activation. A. Gene expression levels of *CCN1* (left) and *CCN2* (right) in PSN1 WT and HPKO cells cultured at low and high confluency. Data is shown as mean□±□SD. **; P<0.01, N.S; not significant, Student’s t-test with Bonferroni correction. B. Immunoblot of PSN1 WT and HPKO cells cultured at low and high confluency. Proteins were detected with indicated antibodies. C. Semi-quantification of band intensities shown in (B). Data is presented as mean□±□SD. *; P<0.05, **: P<0.01, Tukey–Kramer HSD test. D. Immunoblots of PSN1 WT cells at high confluency treated with 100 or 500 nM dasatinib for 2 hours. Proteins were detected with the indicated antibodies. E. Semi-quantification of band intensities shown in (D). Data is shown as mean□±□SD. **; P<0.01, Tukey–Kramer HSD test. F. Immunocytochemistry of PSN1 WT and HPKO cells cultured at low and high confluency. Cells were labeled for YAP (green) and nuclei (DAPI, blue). Scale bar, 50µm. G. Box-and-Whisker plot showing the percentages of cells with nuclear YAP at high confluency in PSN1 WT and HPKO. N=3 for each group, **: P<0.01, Wilcoxon rank sum test. H. Gene expression levels of *CCN1* (left), *CCN2* (middle), and *ANKRD1* (right) in PSN1 WT and HPKO cells cultured at high confluency, with HPKO cells additionally treated with purified proHp (+proHp). Data is shown as mean□±□SD. **; P<0.01, N.S; not significant, Student’s t-test with Bonferroni correction.

To further assess whether extracellular proHp can stimulate expression of YAP target genes, we manipulated proHp levels in the culture environment. Supplementation with purified proHp restored expression of *CCN1* and *ANKRD1* in HPKO cultures (Fig. 5H). Likewise, conditioned medium from PSN1 WT cells, but not HPKO cells, maintained higher expression of *CCN1*, *CCN2*, and *ANKRD1* in PK45 cells (Fig. S4), a non–proHp-producing pancreatic cancer cell line (Fig. 1E). These findings indicate that proHp sustains YAP nuclear localization and target gene expression both at high cell density, thereby allowing pancreatic cancer cells to evade Hippo pathway-mediated growth arrest.

### proHp-induced malignant phenotypes are mediated via YAP activation

We next examined whether changes in YAP activity account for the proHp-dependent growth and motility of pancreatic cancer cells. Forced expression of YAP in PSN1 HPKO cells (hereafter referred to as HPKO+YAP OE) (Fig. S5A) restored the induction of YAP target genes at high confluency, as reflected by increased expression of *CCN1* and *CCN2* genes at high-density conditions (Fig. 6A). In line with this transcriptional rescue, HPKO+YAP OE cells substantially recovered high-confluency proliferation (Fig. 6B). To complement these gain-of-function approaches, the requirement for YAP activity was further tested using pharmacological inhibition. Peptide 17, a peptide inhibitor that blocks the interaction between YAP and the transcription factor TEAD, significantly suppressed the proliferation of PSN1 WT cells at high density, reducing it to the level of HPKO cells (Fig. 6C). In contrast, HPKO cells, which already exhibit attenuated YAP activation at high confluency, showed no additional growth inhibition upon peptide 17 treatment (Fig. 6C), indicating that YAP inhibition phenocopied the loss of proHp.

**Figure. 6.**
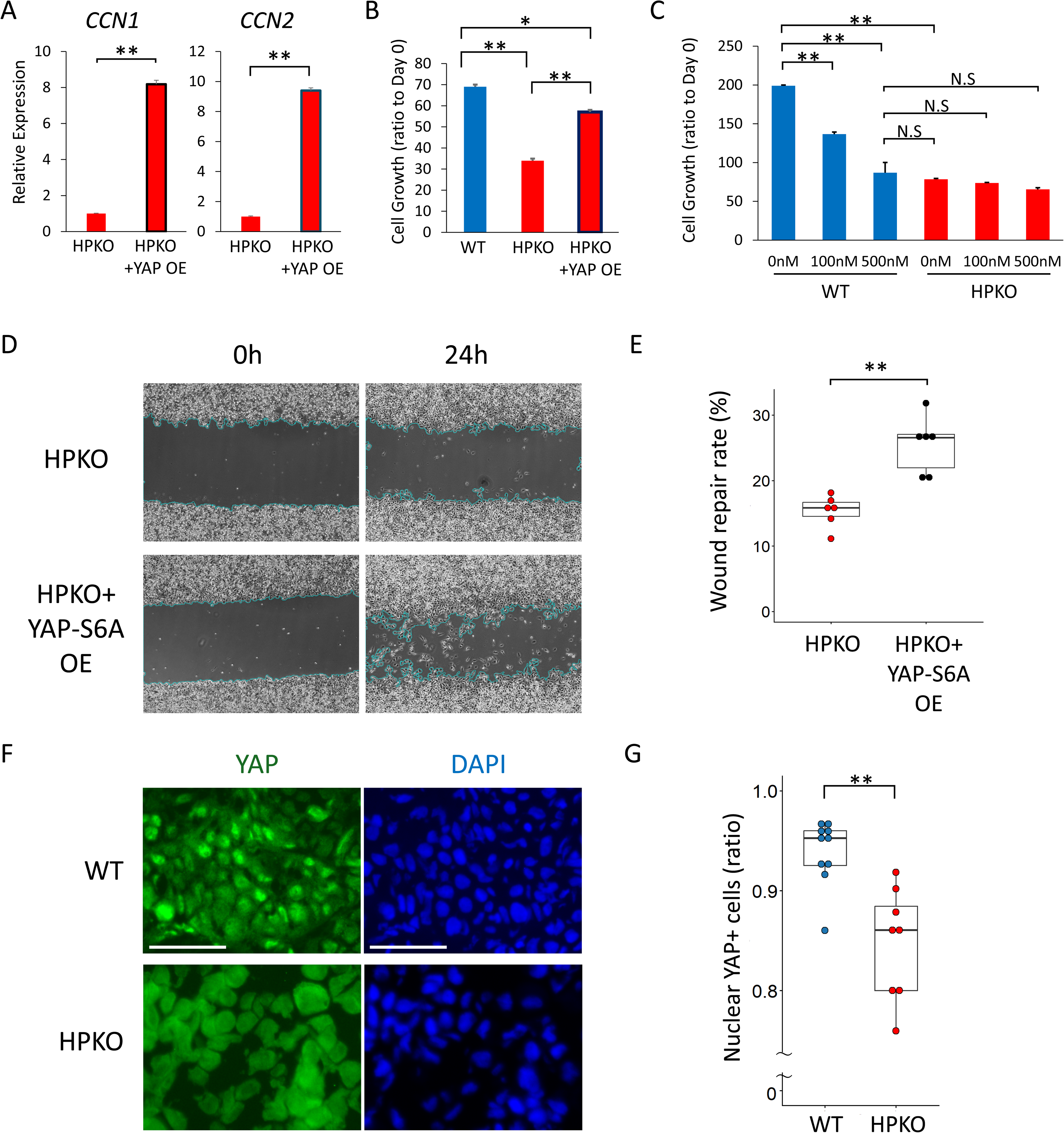
proHp-induced malignant phenotypes are mediated via YAP activation. A. Gene expression levels of *CCN1* (left) and *CCN2* (right) in PSN1 HPKO cells transduced with mock or YAP (+YAP OE), cultured beyond confluence. Data is shown as mean□±□SD. **: P<0.01, Student’s t-test with Bonferroni correction. B. Cell growth assay of PSN1 WT and HPKO cells transduced with mock or YAP (+YAP OE), cultured beyond confluence. Viability was normalized to that at day 0. Data is shown as mean□+□SD. *: P<0.05, **: P<0.01, Student’s t-test with Bonferroni correction. C. Cell growth assay of PSN1 WT and HPKO cells treated with peptide17, a YAP inhibitor, cultured beyond confluence. Viability was normalized to that at day 0. Data is shown as mean□+□SD. **: P<0.01, N.S; not significant, Student’s t-test with Bonferroni correction. D-E. Representative phase-contrast images of wound repair assays for PSN1 HPKO cells transduced with mock or with YAP-S6A (+YAP-S6A OE) at 0 and 24 hours after scratching (D). Box-and-whisker plots quantifying wound repair rates are shown in (E). **; P<0.01, N.S; Not Significant, Wilcoxon rank sum test. F-G: Immunohistochemistry of representative PSN1 WT and HPKO xenograft tumors (F). Cells were labeled for YAP (green) and nuclei (DAPI, blue). Scale bar, 50µm. Boxlzlandlzlwhisker plot showing the percentages of cells with nuclear YAP in PSN1 WT and HPKO tumors (G). Each group consisted of three tumors, and two to three fields per tumor were analyzed. **: P<0.01, Wilcoxon rank sum test.

We also assessed whether the proHp-dependent enhancement of cell motility is mediated via YAP. Although HPKO+YAP OE cells showed improved wound repair in terms of crude repaired area, the wound repair rate did not reach statistical significance (Fig. S5B, C). We therefore introduced a phosphorylation-resistant YAP mutant (YAP-S6A), in which six serine residues targeted by the canonical Hippo pathway are substituted with alanine, into HPKO cells (HPKO+YAP-S6A OE; Fig. S5D). In this setting, wound repair was significantly restored (Fig. 6D, E), indicating that inhibition of Hippo-mediated phosphorylation and degradation of YAP can rescue the motility defect caused by proHp loss.

We further examined YAP behavior *in vivo* under proHp-sufficient and -deficient conditions. Immunohistochemical analysis of subcutaneous PSN1 tumors revealed prominent nuclear localization of YAP in WT tumors, whereas HPKO tumors showed a significant reduction in nuclear YAP (Fig. 6F, G). Taken together, these findings demonstrate that the proHp-dependent promotion of motility and sustained proliferation, both in vitro and in vivo, is mediated through YAP, establishing YAP as a critical downstream effector of proHp in pancreatic cancer progression.

## Discussion

In this study, we demonstrate that prohaptoglobin (proHp) promotes malignant phenotypes of pancreatic cancer and uncover a previously unrecognized link between proHp and YAP activation. Serum proHp levels were elevated in patients with pancreatic cancer compared with healthy controls. Phenotypic analyses showed that proHp enhances cell motility, proliferation under high-density conditions, and tumor growth in xenograft models. Furthermore, transcriptomic and functional analyses indicated that YAP activation is required for the proHp-associated proliferative advantage, implicating the Hippo/YAP pathway as a key downstream effector of proHp signaling in pancreatic cancer cells.

The origin of elevated serum proHp in patients with pancreatic cancer likely involves multiple sources. Hp is predominantly produced in hepatocytes, where proHp is efficiently cleaved by the liver-enriched protease C1RL into mature Hp under physiological conditions ^15^. In contrast, our current data show that a subset of pancreatic cancer tissues exhibits high *HP* expression and that certain pancreatic cancer cell lines secrete proHp, suggesting that tumor tissue can contribute to circulating proHp in addition to the liver. Tumor-infiltrating immune cells can also produce proHp ^31^, and insufficient C1RL activity in these extrahepatic sources may impair complete maturation of proHp. This scenario is reminiscent of procalcitonin, a precursor of the thyroid hormone calcitonin: upon severe microbial infection ^35^ and several other conditions including cancer ^36^, non-thyroidal tissues produce procalcitonin in response to cytokines such as interleukin-6 (IL-6), tumour necrosis factor-α (TNF-α) and interleukin-1β (IL-β) ^37^. Because these tissues lack the specific processing enzymes, procalcitonin is secreted in its precursor form ^37^. A similar mechanism may underlie the accumulation of proHp in the circulation of patients with pancreatic cancer. Regardless of the precise cellular origin, our functional studies indicate that elevated proHp can actively promote tumor progression, suggesting that proHp is not merely a passive marker of systemic stress but as a context-dependent modulator of pancreatic cancer advancement.

The Hippo signaling pathway is a central regulator of cell proliferation and organ size ^38^. When activated (Hippo-on state), LATS1/2-mediated phosphorylation of the transcriptional co-activators YAP/TAZ promotes their cytoplasmic sequestration and degradation, thereby suppressing their transcriptional activity. Conversely, when inactive (Hippo-off state), dephosphorylated YAP/TAZ translocate to the nucleus and activate target genes expression ^32, 33, 38^. In cancer, inappropriate YAP/TAZ activation drives malignant progression, including resistance to contact-mediated growth inhibition ^32, 33^, processes that are directly relevant to the context-dependent phenotypes we observed in pancreatic cancer cells with either endogenous or exogenous proHp.

Our RNA-sequencing analysis showed that proHp is associated with maintenance of YAP target gene expression under high-confluency conditions, where these genes are normally suppressed. While HPKO cells showed a marked reduction in target expression upon confluence, WT cells maintained their expression, suggesting that proHp blunts full contact inhibition and preserves YAP-dependent transcription at high cell density. Functional experiments were consistent with this model: YAP overexpression partially restored high-density proliferation in HPKO cells, whereas pharmacological inhibition of YAP-TEAD interaction phenocopied the growth inhibition upon depletion of proHp. The partial rescue by YAP overexpression suggests that while YAP is a central effector of proHp, additional pathways may also contribute. Taken together, these data indicate that proHp promotes pancreatic cancer progression at least in part by maintaining YAP activity and attenuating Hippo pathway–mediated growth arrest under high-density conditions.

Unlike many signal pathways that depend on specific ligand-receptor pairs, the Hippo pathway integrates diverse physical and biochemical inputs, including cell-cell contact, extracellular matrix (ECM) mechanics, and soluble cues ^32, 38^. Cells can sense mechanical properties of their environment—such as stiffness and deformation—and transmit these signals through mechanotransducers, cytoskeletal tension sensors, and adhesion complexes. Consequently, Hippo pathway can be modulated without relying on a specific ligand-receptor pair, in contrast to many other signaling cascade ^33^. GPCRs, integrins, and cell-cell adhesion molecules converge on Hippo regulation, and pancreatic cancer tissue is characterized by a dense, stiff stroma that promotes YAP/TAZ-mediated transcription ^39^. Given that mature Hp has been reported to modulate ECM integrity through inhibition of gelatinases such as MMP-2 and MMP-9 ^40^, it is conceivable that proHp may also influence ECM organization and thereby indirectly support YAP activation. SRC family kinases occupy nodal positions downstream of integrins and adhesion complexes and are key components of mechanotransduction pathways that impinge on YAP. Consistent with this, noncanonical regulation of YAP through SRC family kinase (SFK)–mediated phosphorylation at Y357 has been reported to promote YAP nuclear localization independently of S127 status ^34^. In our experiments, high-density WT PSN1 cultures showed SFK activation and increased Y357 phosphorylation of YAP that were sensitive to the SFK inhibitor dasatinib. Although these data do not establish a direct causal link between Y357 phosphorylation and the phenotypes observed, they raise the possibility that proHp engages SFK-dependent signaling that contributes to maintenance of nuclear YAP at high cell density.

Taken together, this study identifies proHp as a context-dependent promoter of pancreatic cancer progression that acts, at least in parts, by sustaining YAP activity and attenuating Hippo pathway-mediated growth control in response to the mechanical microenvironment. By linking a secreted proHp to YAP-dependent growth control within the pancreatic tumor microenvironment, our findings suggest that proHp can modulate malignant behavior under specific cellular and tissue contexts. Because both the responsiveness to proHp and its post-translational modification state are likely to influence these contexts, further investigations are required to determine how the proHp-YAP axis can be used to stratify patients. Nevertheless, our results define the proHp-YAP axis as a tractable mechanism for future investigation of biomarkers and therapeutic targets in pancreatic cancer.

## Data Availability

RNA-seq data are available at DNA Data Bank of Japan (DDBJ) Sequence Read Archive (DRA) under accession number DRA028777. The other data are available from the corresponding authors upon reasonable request.

## Acknowledgements

We acknowledge the NGS core facility at the Research Institute for Microbial Diseases of The University of Osaka for the sequencing and data analysis. The authors used an AI-based language model (Perplexity, powered by GPT-5.1) to assist in editing and polishing the English of this manuscript. The authors take full responsibility for the content of the manuscript.

## CRediT Authorship Contributions

Jumpei Kondo (Conceptualization: Lead; Methodology: Supporting; Formal analysis: Lead; Data curation; Lead; Visualization: Lead; Writing – original draft: Lead; Writing – review & editing: Lead)

Honoka Nakayama (Investigation: Lead; Methodology: Equal; Formal analysis: Supporting; Visualization: Supporting; Writing – review & editing: Supporting)

Ayusa Kuroda (Investigation: Lead; Methodology: Equal; Formal analysis: Supporting; Visualization: Supporting; Writing – review & editing: Supporting)

Ayumu Hayashibara (Investigation: Lead; Methodology: Equal; Formal analysis: Supporting; Visualization: Supporting; Writing – review & editing: Supporting)

Daisuke Sakon (Conceptualization: Supporting; Data curation: Supporting;

Methodology: Supporting; Investigation: Supporting; Visualization: Supporting; Writing – original draft: Supporting; Writing – review & editing – Equal)

Shinji Takamatsu (Supervision: Lead; Methodology: Supporting; Resources: Supporting; Writing – review & editing: Supporting)

Hirofumi Akita (Resources: Lead; Investigation: Supporting; Writing – review & editing: Supporting)

Hidetoshi Eguchi (Resources: Lead; Investigation: Supporting; Writing – review & editing: Supporting)

Eiji Miyoshi (Conceptualization: Supporting; Supervision: Lead: Project administration: Lead; Funding acquisition: Lead; Writing – review & editing: Equal)

## Abbreviations

ANOVA: (analysis of variance)
DAPI: (4′,6-diamidino-2-phenylindole)
ECM: (extracellular matrix)
ELISA: (enzyme-linked immunosorbent assay)
EMT: (epithelial–mesenchymal transition)
FBS: (fetal bovine serum)
FDR: (false discovery rate)
FFPE: (formalin-fixed paraffin-embedded)
FPKM: (fragments per kilobase of exon per million mapped reads)
Fuc-Hp: (fucosylated haptoglobin)
GO: (Gene Ontology)
GPCR: (G protein–coupled receptor)
H&E: (hematoxylin and eosin)
Hb: (hemoglobin)
Hp: (haptoglobin)
HPKO: (HP-knockout)
HV: (healthy volunteer)
IL: (interleukin)
KO: (knockout)
MMP: (matrix metalloproteinase)
NES: (normalized enrichment score)
NBF: (neutral buffered formalin)
N.S.: (not significant)
OE: (overexpression)
PC: (pancreatic cancer)
PBS: (phosphate-buffered saline)
PDAC: (pancreatic ductal adenocarcinoma)
PEI: (polyethyleneimine)
PFA: (paraformaldehyde)
proHp: (prohaptoglobin)
PVDF: (polyvinylidene difluoride)
qPCR: (quantitative polymerase chain reaction)
RIPA: (radioimmunoprecipitation assay)
RNA-seq: (RNA sequencing)
RT-PCR: (reverse transcription polymerase chain reaction)
SD: (standard deviation)
SFK: (Src family kinase)
TAZ: (transcriptional co-activator with PDZ-binding motif)
TBST: (Tris-buffered saline with Tween 20)
TEAD: (TEA domain family member)
TNF: (tumor necrosis factor)
VEGF: (vascular endothelial growth factor)
WT: (wild-type)
YAP: (Yes-associated protein)
GSEA: (gene set enrichment analysis)

