## Supplementary material for "Prohaptoglobin Promotes Pancreatic Cancer Progression by sustaining YAP Activity": Fig S

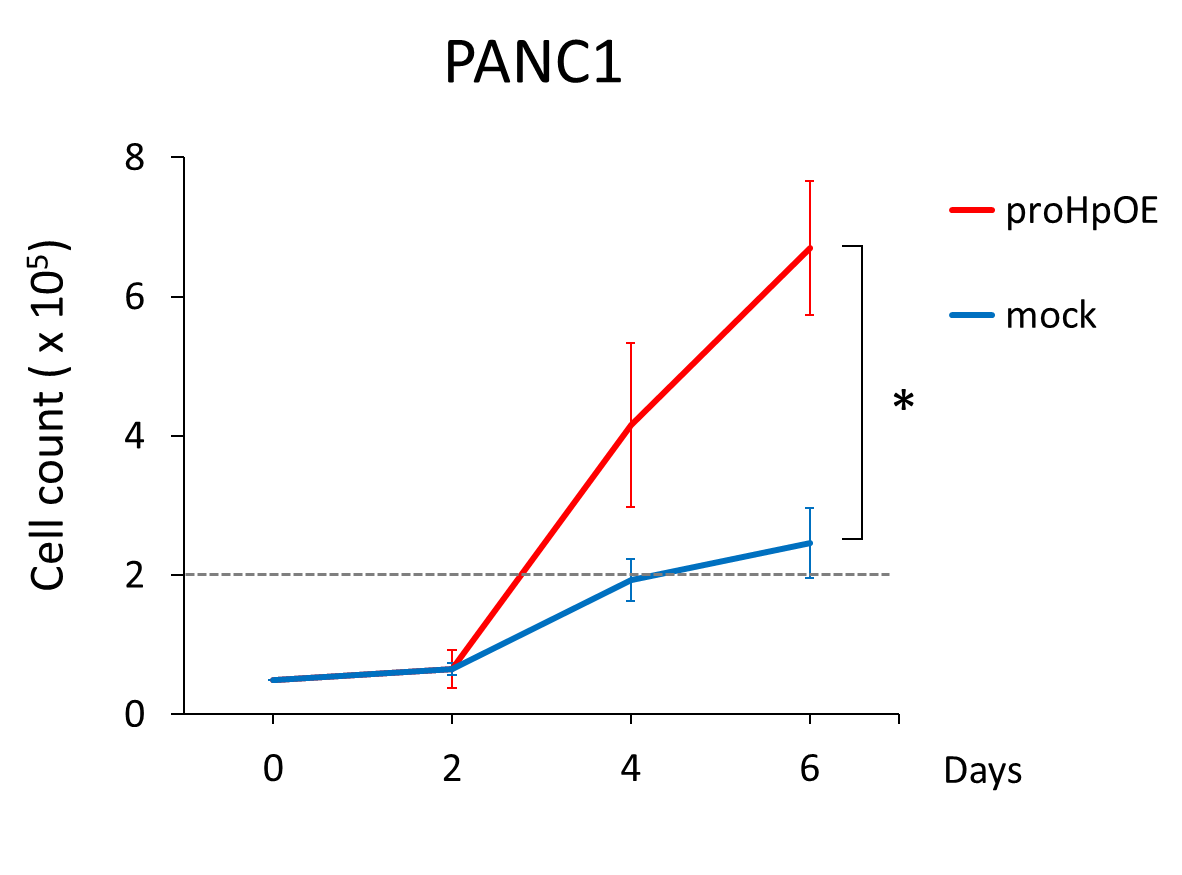

**Figure. S1 Prohaptoglobin promotes long-term proliferation of PANC1 cells beyond confluence**

Growth curves of PANC-1 mock-transfected (mock, blue) and prohaptoglobin-overexpressing (proHpOE, red) cells cultured for an extended period. The horizontal dotted line denotes the cell density at which cells are visually confluent. Data are shown as mean ± SD for N = 3 per condition. *P < 0.05, repeated-measures ANOVA.

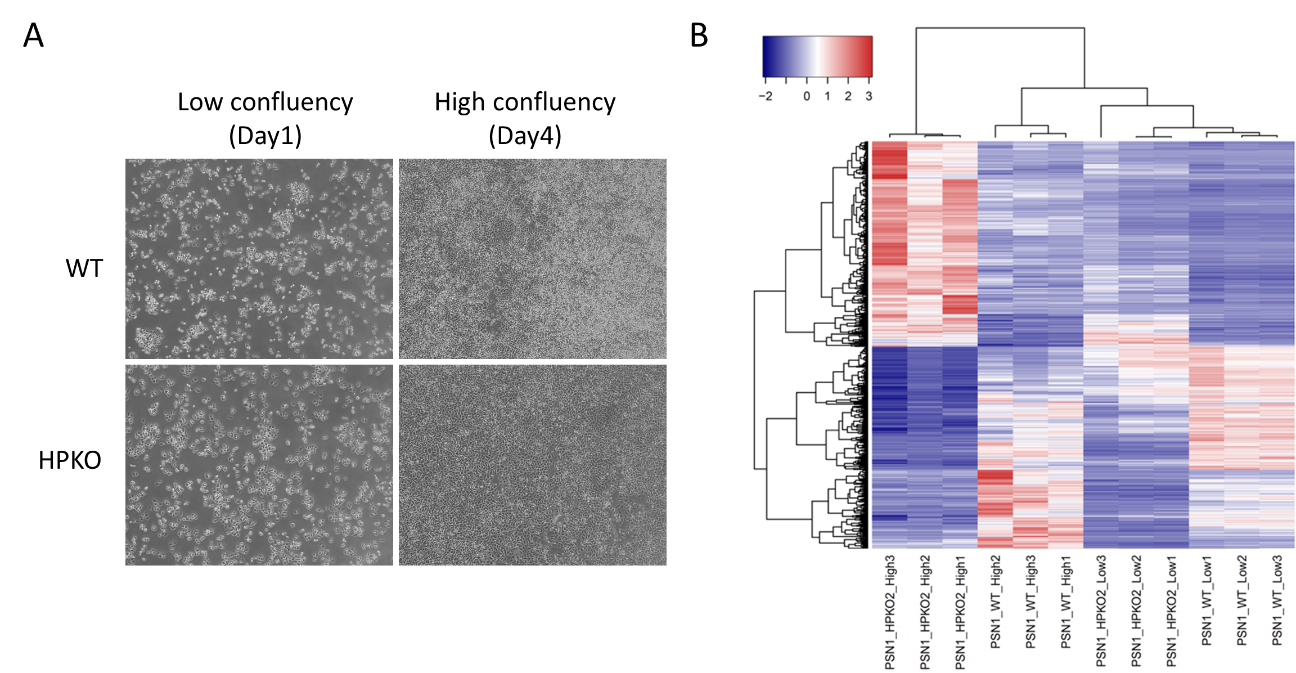

Figure. S2 Transcriptomic overview of PSN1 WT and HPKO cells at low and high confluency.

A. Representative phase-contrast images of PSN1 wild-type (WT) and HP knockout (HPKO) cells cultured at low and high confluency, corresponding to the time points used for RNA-seq sample collection.

B. Heatmap of differentially expressed genes (DEGs) in PSN1 WT and HPKO cells at low and high confluency.

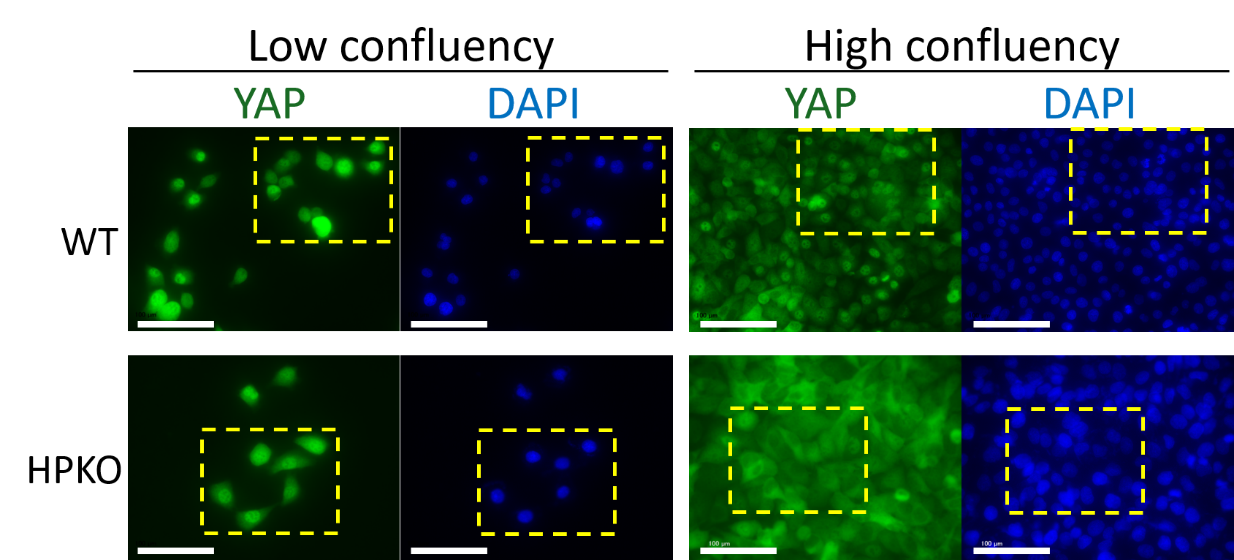

Figure. S3 Subcellular distribution and immunofluorescence of YAP in pancreatic cancer cells

Immunocytochemistry of PSN1 WT and HPKO cells cultured at low and high confluency. Cells were labeled for YAP (green) and nuclei (DAPI, blue). Scale bar, 100µm. Yellow dotted-boxed areas are enlarged in Figure 5E.

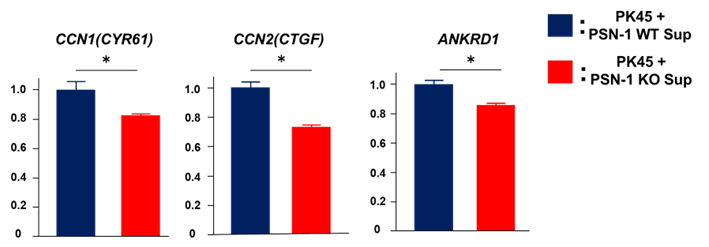

Figure. S4 Prohaptoglobin-containing conditioned medium sustains YAP target gene expression in PK45 cells

Gene expression levels of *CCN1* (left), *CCN2* (middle), and *ANKRD1* (right) in PK45 cells cultured at high confluency and treated with culture supernatant from PSN1 WT or HPKO cells. Data are shown as mean ± SD. *P < 0.05, Student’s t-test with Bonferroni correction.

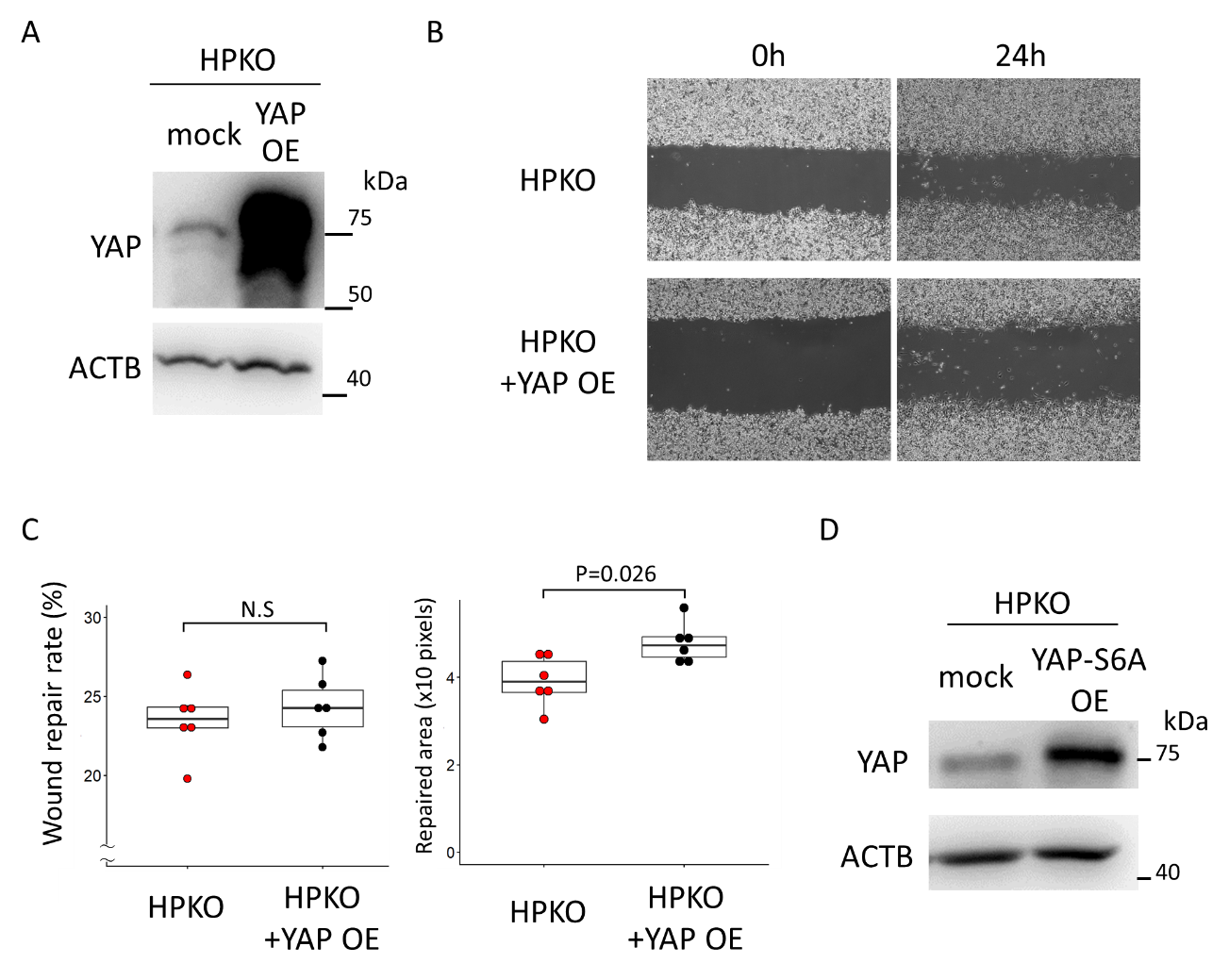

Figure. S5 YAP and YAP-S6A overexpression in PSN1 HPKO cells

A. Immunoblot analysis of PSN1 HPKO cells transduced with mock or YAP-overexpressing (YAP OE) lentiviral vectors. Proteins were detected with the indicated antibodies.

B-C. Wound repair assays for PSN1 HPKO cells. Representative phase-contrast images of PSN1 HPKO cells transduced with mock or YAP OE at 0 and 24 hours after scratching are shown in (B). Box-and-whisker plots of wound repair rate (C, left) and crude repaired area (C, right) are shown. N.S; Not Significant, Wilcoxon rank sum test.

D. Immunoblot analysis of PSN1 HPKO cells transduced with mock or YAP- overexpressing (YAP-S6A OE) lentiviral vectors. Proteins were detected with the indicated antibodies.
